# Human stem cell resources are an inroad to Neandertal DNA functions

**DOI:** 10.1101/309658

**Authors:** Michael Dannemann, Benjamin Vernot, Svante Pääbo, Janet Kelso, J. Gray Camp

## Abstract

Pluripotent stem cells from diverse humans offer the potential to study human functional variation in controlled culture environments. A portion of this variation originates from ancient admixture between modern humans and Neandertals, which introduced alleles that left a phenotypic legacy on individual humans today. Here we show that a large repository of human induced pluripotent stem cells (iPSCs) harbors extensive Neandertal DNA, including most known functionally relevant Neandertal alleles present in modern humans. This resource contains Neandertal DNA that contributes to human phenotypes and diseases, encodes hundreds of amino acid changes, and alters gene expression in specific tissues. Human iPSCs thus provide an opportunity to experimentally explore the Neandertal contribution to present-day phenotypes, and potentially study Neandertal traits.

## MAIN TEXT

Protocols have been developed to differentiate human embryonic and induced pluripotent stem cells (iPSCs) into many different cell types of the human body^1^. In addition, stem cells can self-organize into complex three-dimensional structures containing multiple cell types that resemble human tissues (such as the brain, liver, stomach, intestine, and kidney)^2^. These stem cell-derived systems can be used to explore how natural variation between human individuals impacts development and cell biology^3^. Some of the variation in present-day humans derives from admixture between modern and archaic hominids. Analyses of Neandertal genomes have shown that Neandertals and modern humans interbred approximately 55,000 years ago as the latter migrated out of Africa. As a consequence, around 2 percent of the genomes of all present-day non-Africans derive from Neandertal ancestors^4–6^. Because the segments of DNA inherited from Neandertals varies between individuals, cumulatively at least 40% of the Neandertal genome survives in people today^7^. Recent genome-wide association studies suggest that the DNA relics from this admixture left an extensive phenotypic legacy, influencing for example skin and hair color, immune response, lipid metabolism, bone morphology, blood coagulation, sleep patterns, and mood disorders^7–15^. In addition, Neandertal-derived DNA has a significant effect on gene expression in adult human tissues^16,17^. However, these associations have been observed in living people or in tissues, where there is limited opportunity for controlled experimentation. Further, there are few opportunities to study the impact of Neandertal-derived DNA on modern human development. Recently, the Human Induced Pluripotent Stem Cell Initiative (HipSci) published their work on generating and characterizing a large resource of human iPSCs with genome-wide genotype data^18^. This large repository presents an unprecedented opportunity to identify carriers of Neandertal alleles of interest and explore the genetic mechanisms underlying Neandertal and modern human phenotypes.

We have analyzed the genome sequences from 173 of these individuals (mostly Europeans) and identified the modern-human and Neandertal component of each individual’s ancestry (Fig. 1A, S1). We used alleles in present-day humans that are shared with the Vindija Neandertal and absent in Yoruba individuals, along with a linkage disequilibrium-based test for incomplete lineage sorting (ILS), to identify haplotypes that are likely of Neandertal origin (Fig. S1B-C). We used the Vindija Neandertal genome to identify Neandertal haplotypes because it is genetically more similar to the introgressing Neandertals than the Altai individual^5^, thus providing additional power to detect haplotypes^6^. Based on these inferred haplotypes, we find that cumulatively 19.6% (661 Mb) of the Neandertal genome is represented in these cell lines, with between 18.7 and 30.9 Mb Neandertal DNA per individual (Table 1, S1–S3). We found that 98% of inferred haplotypes overlap previously identified introgressed sequence and that the cumulative amount of Neandertal DNA present in this resource approaches the total amount that has been identified in Europeans^19^ (21.3%; Fig. 1B). We note that the addition of non-European populations would extend this even further. For example, an additional 16.4% of the Neandertal genome has been identified in east and south Asians^19^, but is absent from the HipSci resource as individuals are largely of European ancestry.

**Figure 1:**
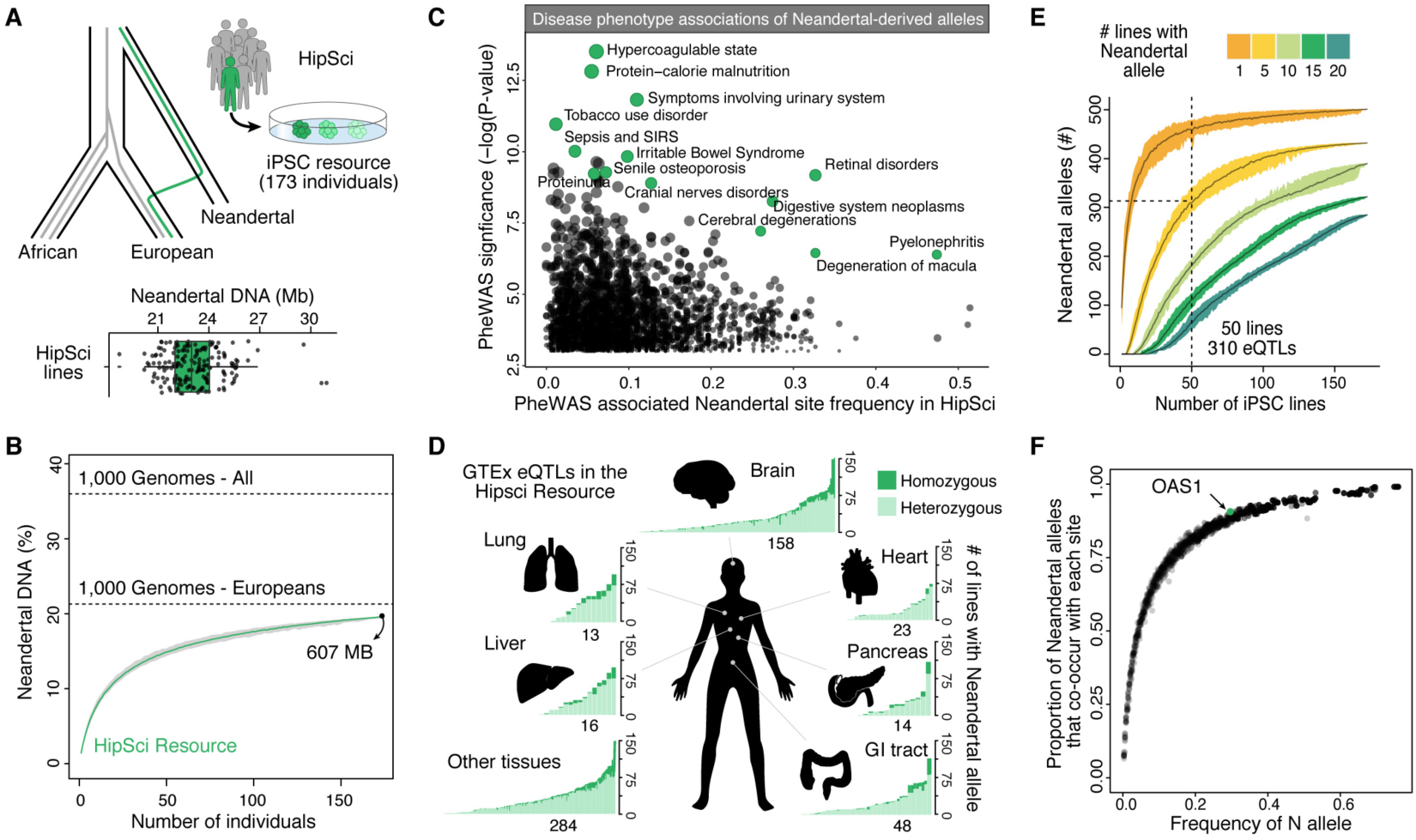
The HipSci resource harbors extensive functionally relevant Neandertal DNA. (A) The Human Pluripotent Stem Cell Initiative (HipSci) created and characterized induced pluripotent stem cells lines from 173 individuals with genome-wide genotype data, which we analyzed for Neandertal ancestry. The boxplot shows Neandertal DNA in megabases (Mb) per individual. (B) The cumulative percentage of the human genome covered by Neandertal haplotypes in the HipSci resource. The percentage is also shown for European and all non-African individuals from the 1000 genome project. (C) The frequency of single nucleotide changes introgressed from Neandertals that have significant associations with human disease phenotypes^8^, with select associations highlighted in teal. (D) The number of iPSC lines that contain each Neandertal eQTL from the GTEx dataset present in the HipSci resource, colored by tissue and by homozygous (dark) or heterozygous (light). (E) A power analysis shows how many Neandertal eQTLs are present in at least a certain number of cell lines (color) out of a random sample of X lines (x-axis) from the HipSci resource. (F) Plot showing the co-occurrence of Neandertal alleles within individuals. The frequency of each Neandertal allele is plotted against the proportion of all other Neandertal alleles with which it co-occurs in the HipSci resource. For example, an OAS1 Neandertal-derived allele is found at relatively high frequency (0.29, and present in 53% of all individuals) in the HipSci resource and is therefore paired with the majority of other Neandertal alleles.

**Table 1:** Breakdown of Neandertal DNA coverage in the HipSci resource.

|  | Average cell lines | Cumulative cell lines |
| --- | --- | --- |
| Neandertal ancestry (bps) | 22,985,635 (18,725,364-30,931,585) | 607,404,493 |
| Haplotype length (bps) | 48,014.5 (42,613-56,968) | - |
| Neandertal alleles | 15,844.9 (12,829-20,060) | 205,032 |
| Missense alleles | 58.6 (38-90) | 719 |
| Alleles in enhancers/promoters | 235.4 (169-340) | 2,852 |
| PheWAS alleles | 1.5 (0-5) | 924 |
| GTEx eQTLs | 44.6 (29-60) | 270 |
| Allele-specific expression | 119.3 (75-176) | 957 |

We next analyzed the power of the HipSci resource to study functionally relevant Neandertal DNA. We collected recently published Neandertal-derived phenotype and disease associated alleles^8,10–14^ and found that most (22/24) of the alleles that reached genome-wide significance are present in the resource in more than 1 iPSC line (Table 2). These alleles are associated with a variety of processes including digestive function, nutrition, skin color, coagulatory protein production, and immune response (Fig. 1C). In addition, we identified hundreds of alleles that alter amino acids, are expression quantitative trait loci (eQTL)^14^ or show allele-specific expression^16^ (Fig. 1D; Table S1).

**Table 2:**
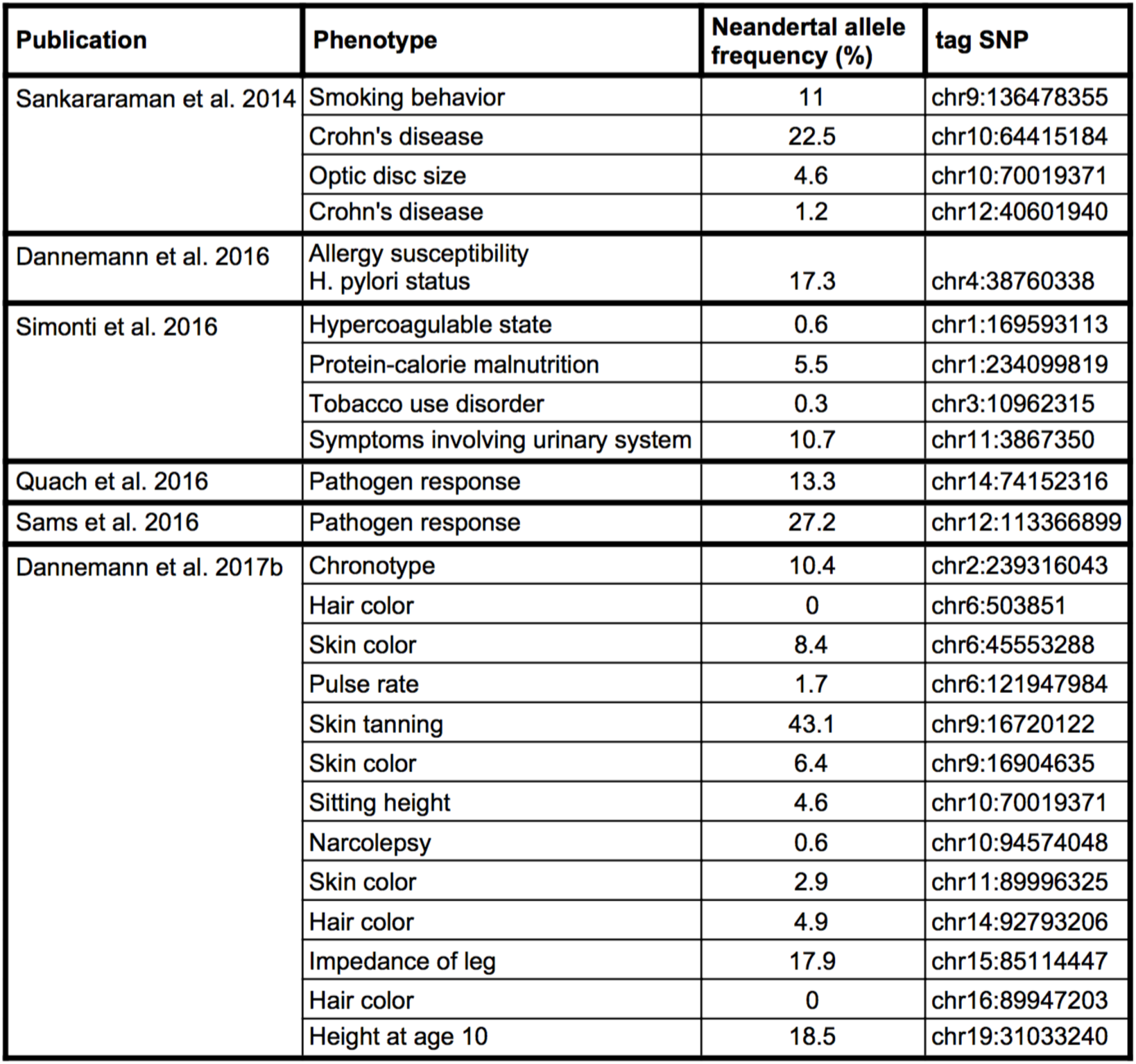
Prevalence of phenotype associated Neandertal alleles in the HipSci resource.

| Publication | Phenotype | Neandertal allele frequency (%) | tag SNP |
| --- | --- | --- | --- |
| Sankararaman et al. 2014 | Smoking behavior | 11 | chr9:136478355 |
|  | Crohn's disease | 22.5 | chr10:64415184 |
|  | Optic disc size | 4.6 | chr10:70019371 |
|  | Crohn's disease | 1.2 | chr12:40601940 |
| Dannemann et al. 2016 | Allergy susceptibility<br>H. pylori status | 17.3 | chr4:38760338 |
| Simonti et al. 2016 | Hypercoagulable state | 0.6 | chr1:169593113 |
|  | Protein-calorie malnutrition | 5.5 | chr1:234099819 |
|  | Tobacco use disorder | 0.3 | chr3:10962315 |
|  | Symptoms involving urinary system | 10.7 | chr11:3867350 |
| Quach et al. 2016 | Pathogen response | 13.3 | chr14:74152316 |
| Sams et al. 2016 | Pathogen response | 27.2 | chr12:113366899 |
| Dannemann et al. 2017b | Chronotype | 10.4 | chr2:239316043 |
|  | Hair color | 0 | chr6:503851 |
|  | Skin color | 8.4 | chr6:45553288 |
|  | Pulse rate | 1.7 | chr6:121947984 |
|  | Skin tanning | 43.1 | chr9:16720122 |
|  | Skin color | 6.4 | chr9:16904635 |
|  | Sitting height | 4.6 | chr10:70019371 |
|  | Narcolepsy | 0.6 | chr10:94574048 |
|  | Skin color | 2.9 | chr11:89996325 |
|  | Hair color | 4.9 | chr14:92793206 |
|  | Impedance of leg | 17.9 | chr15:85114447 |
|  | Hair color | 0 | chr16:89947203 |
|  | Height at age 10 | 18.5 | chr19:31033240 |

We find that 50 HipSci lines chosen at random will allow the interrogation of ~310 Neandertal-associated eQTLs with each site represented in at least 5 cell lines (Fig. 1E, S2). We note that each Neandertal allele present in the HipSci resource exists in a primarily modern human background. However, in each individual many Neandertal alleles co-occur (Fig. 1F). For example, in the HipSci resource, one of the Neandertal alleles at the OAS1 locus (chr12:113425154), a locus with the highest Neandertal frequencies in present-day humans, is paired with 90% of the other introgressed Neandertal alleles in at least one cell line. It may thus be possible to leverage such co-occurrences to study epistatic interactions among Neandertal alleles. Additionally, as iPSC resources continue to expand to include individuals from other populations, it will become possible to explore the phenotypic contribution of alleles derived from other archaic hominins such as Denisovans^20^, a distant Asian relative of Neandertals that made even larger genetic contributions to present-day people in parts of Oceania^21–23^.

Taken together, our analysis suggests that human-Neandertal hybrid iPSC resources can be used to systematically explore Neandertal allele function in diverse cell types differentiated in controlled culture environments, including the previously unexplored study of developmental processes.

## METHODS

### Detection of Neandertal haplotypes

To define Neandertal haplotypes, we first identified a set of SNPs where one allele is likely of Neandertal origin. These Neandertal SNPs (aSNPs) have one allele that is (iii) present in the genomes of the Vindija Neandertal^6^, and (ii) present 1,000 Genomes Project (phase III) Eurasian populations, but (iii) absent from Yoruban, an African population with little to no Neandertal admixture^19^. To detect putative Neandertal haplotypes we scanned for consecutive stretches of aSNPs in the genomes of the cell lines where the individual carries the Neandertal-shared alleles, with continuous SNPs located not more than 20,000 bps from one another. To define a Neandertal haplotype we required a consecutive stretch of at least three Neandertal alleles across their corresponding successive aSNPs. Haplotypes are additionally tested for a length that exceeds the expected length for segments of incomplete lineage sorting, based on the algorithm presented by Huerta-Sanchez et al. ^24^ and the age of the divergence to Neandertals of 465,000 years used in ^12^, a conservative estimate of the human mutation rate, (mu=1x10^−8^ per site per generation) and two recombination maps ^25,26^. The resulting P values have been corrected for multiple testing using the Benjamini-Hochberg approach. We included haplotypes with an FDR < 0.05 for ILS for at least one of the recombination maps, or if no recombination map data was available, inferred haplotypes with a length greater than 50kb or at least 10 consecutive aSNPs with an Neandertal allele to our analyses. All inferred haplotypes for each cell line are available in supplementary data tables. We have applied the method to the genotype data for 173 individuals of the HipSci resource and all non-Africans of the 1,000 Genomes project (phase III).

### Detection of Neandertal missense, regulatory and pheWAS variants

We sought to identify putatively functional Neandertal alleles that overlap confidently inferred Neandertal haplotypes (see section “Detection of Neandertal haplotypes”) in the cell lines by detecting those that alter the protein or regulatory sequence of a gene. First, for the detection of Neandertal alleles that modify the protein sequence, we selected all Neandertal alleles within confidently inferred Neandertal haplotypes detected in any cell line and annotated them functionally using the variant effect predictor (VEP, human Ensembl version 73). We selected those alleles that were defined as ‘missense’ by VEP. Second, we annotated Neandertal alleles likely to be involved in gene regulation by overlapping them with three datasets: (i) enhancer and promoter regions provided by the Ensembl Regulatory Build ^27^, (ii) significant eQTLs in the GTEx dataset ^17^ and (iii) allele-specfic expression ^16^. For (i) we identified aSNPs with the Neandertal alleles directly overlapping with a regulatory motif. For (ii) we selected the inferred Neandertal haplotypes with the top 20 most significant eQTLs in each of the 48 GTEx tissues with more than 50 individuals ^28^ and required to have at least one aSNP to be present in a given Neandertal haplotype and iPSC individual, resulting in a total of 409 such Neandertal haplotypes. For (iii) we selected all Neandertal alleles that have been identified to show allele-specific expression (FDR<0.1). Third, we queried the pheWAS database for associations of Neandertal alleles and specific phenotypes in modern humans. We selected all 925 aSNPs with significant phenotype associations detected by Simonti et al.^8^. We further selected multiple additional significant phenotypes associations for Neandertal alleles (Table 2) ^8,l0–14^.

### Power analysis

The ability to study a particular Neandertal variant depends both on its effect size and its frequency within a given sample of individuals or cell lines. We cannot control the effect size, but one can – within reason – control the number of samples they consider. Larger sample sizes allow more variants to be studied, but may offer diminishing returns. To determine the power of studies of certain sample sizes, we considered how many Neandertal variants would be present at particular frequency thresholds, as an effect of sample size. For each category of Neandertal allele (eQTL, amino acid change, etc), we subsampled X cell lines, and counted the number of Neandertal variants present at least Z times, for values of Z = (1, 5, 10, 15, 20). This subsampling was repeated 100 times for all values of X and Z. We plot the average number of Neandertal variants present at a particular rate on the Y axis. Each possible value of Z is given a different color, and the range of values over 100 resamplings is shown as colored confidence intervals. For example, given a sample of 50 random cell lines, 62% of the 501 Neandertal eQTLs in the HipSci resource are present in at least 5 cell lines, and 92% are present in at least 1 cell line.

### Principle Component Analysis on HipSci lines

To infer the genetic relationship between the HipSci individuals and present-day people, we have performed a principal component analysis using polymorphic sites in 1000 Genomes Eurasians ^19^ that show large population differentiation between Europeans and Asians (Fst>0.5). Population differentiation has been calculated based on Fst, using the Weir and Cockerham calculation implemented in vcftools ^29^ and 100 unrelated Asians and Europeans each in the 1000 Genomes panel. The principle components have been computed using SNPs with Fst > 0.5 between Europeans and Asians in the 1000 Genomes. While the HipSci resource contains non-European individuals, almost all of the individuals with genotypes are clustering with Europeans from the 1000 Genomes panel (Fig. 1B).

## SUPPLEMENTARY INFORMATION

**Figure S1:**
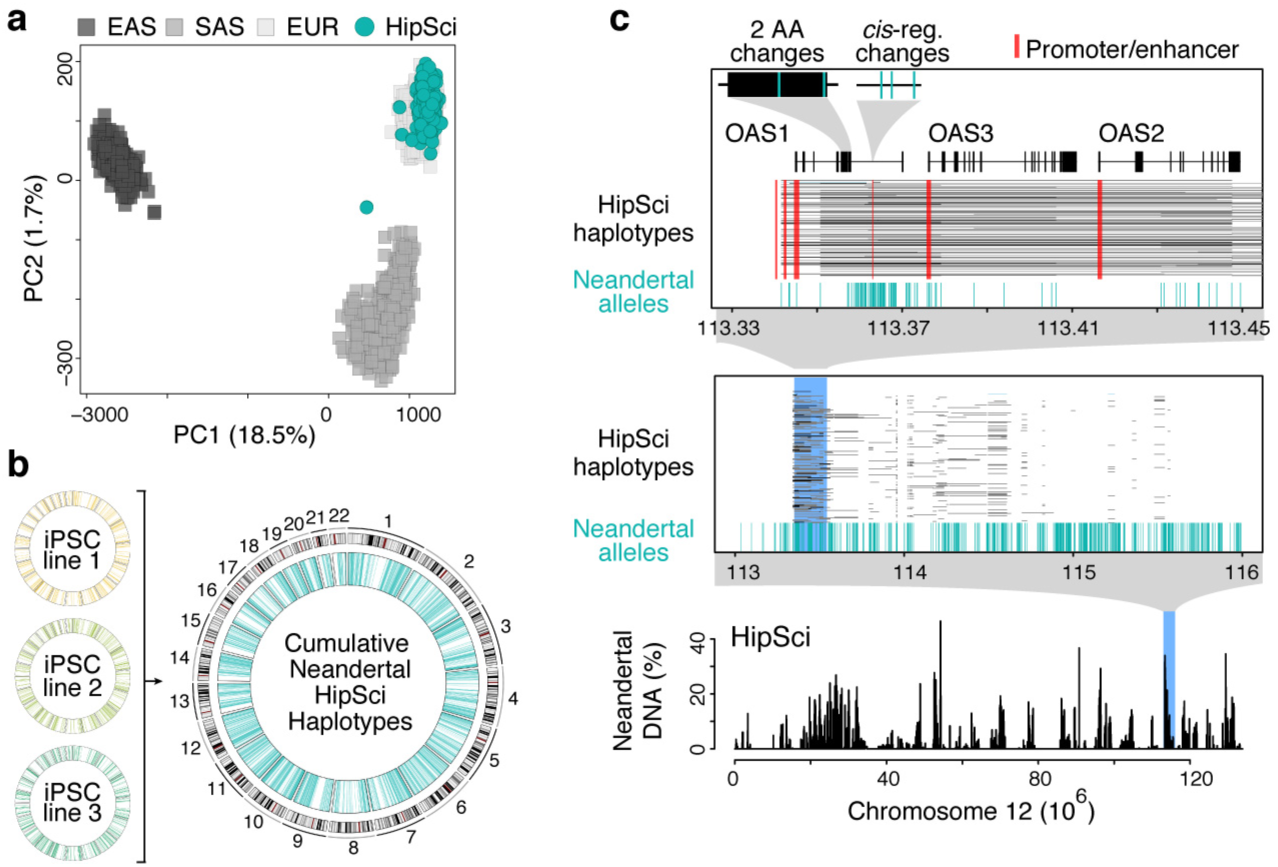
Identification of Neandertal haplotypes in human iPSC lines. (A) Principal component analysis on SNPs that distinguish East Asian (EAS, dark grey), Southeast Asian (SAS, grey), and European (light grey) individuals suggests that each HipSci individual (teal) has a major European component to their ancestry. (B) Neandertal haplotype coverage across each chromosome for three individuals, as well as the cumulative coverage for all 173 HipSci individuals (Circos plots). (C) Neandertal DNA percentage, alleles, and haplotypes present in the HipSci resource covering the OAS locus. In this example, the gene OAS1 contains both regulatory and amino acid changing Neandertal alleles, present at a frequency of approximately 27% in HipSci individuals.

**Figure S2:**
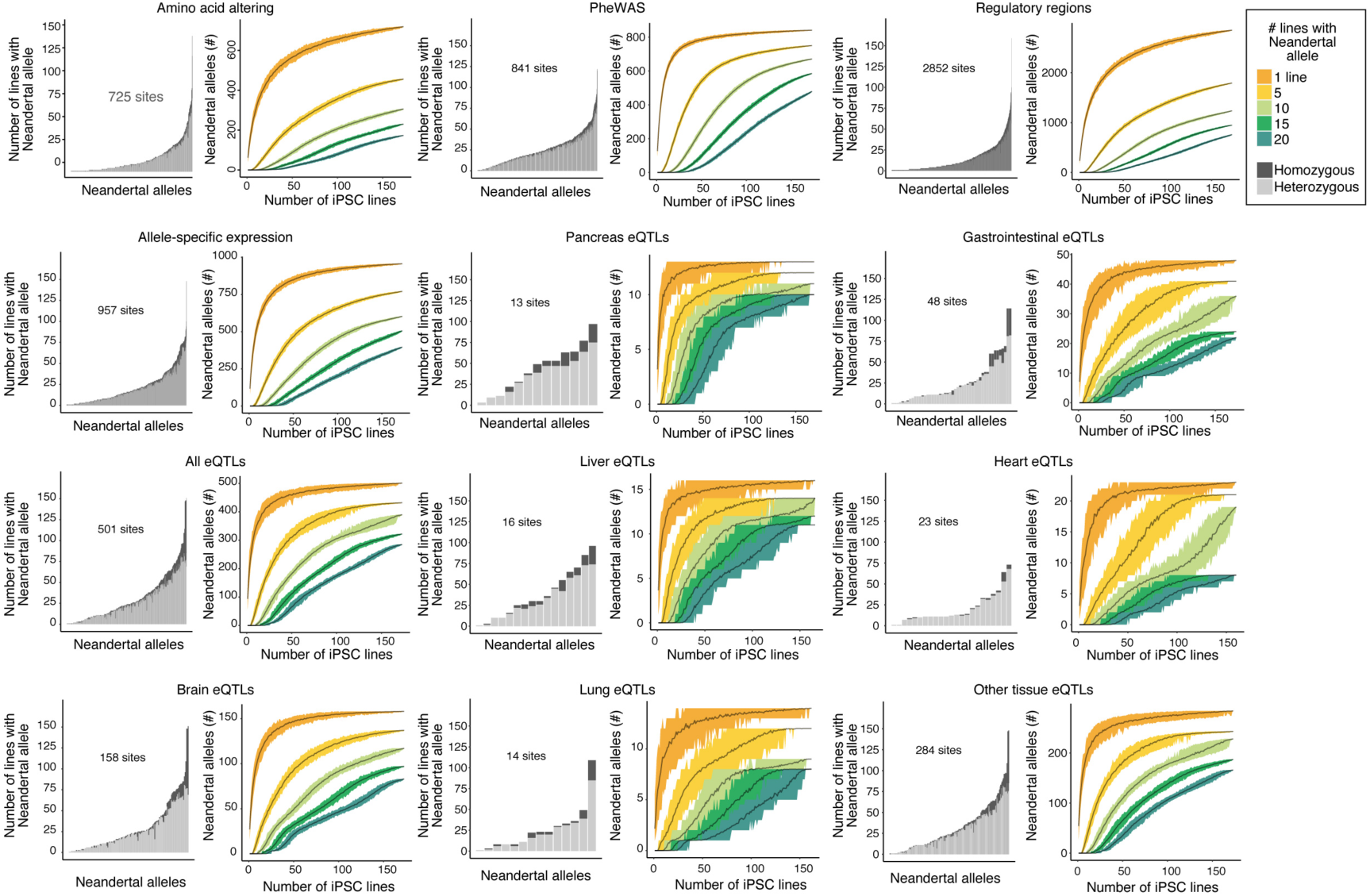
Power analyses for multiple classes of functional alleles. The number of iPSC lines that contain each introgressed allele is shown for multiple different categories of functional alleles, colored by homozygosity (dark) or heterozygosity (light) (left). For each category, a power analysis shows how many introgressed alleles are present in a random sample of X lines (x-axis) (right).

**Table S1:** Neandertal ancestry estimates and functional alleles in HipSci resource.

**Table S2:** Inferred Neandertal haplotypes in HipSci resource based on Vindija Neandertal. Tables for each HipSci individual with all inferred haplotypes including genomic locations, FDRs for tests of compatibility of incomplete lineage sorting and inferred Neandertal alleles (with chromosomal location, hg19 and status of Neandertal allele; 0|1: heterozygous, 1|1: homozygous).

**Table S3:** Inferred Neandertal haplotypes in HipSci resource based on Altai Neandertal. Tables for each HipSci individual with all inferred haplotypes including genomic locations, FDRs for tests of compatibility of incomplete lineage sorting and inferred Neandertal alleles (with chromosomal location, hg19 and status of Neandertal allele; 0| 1: heterozygous, 1|1: homozygous).

## AUTHOR CONTRIBUTIONS

MD, BV, JGC analyzed the data and wrote the manuscript with input from JK and SP.

## AUTHOR INFORMATION

The authors declare no conflicts of interest. Correspondence and requests for materials should be addressed to.

## ACKNOWLEDGEMENTS

This work was supported by the NOMIS Foundation and the Max Planck Society. The GTEx data used for the analyses described in this manuscript were obtained from dbGaP accession number phs000424.v6.p1.c1 on 05/23/2016.

## REFERENCES

1. Williams, L. A., Davis-Dusenbery, B. N. & Eggan, K. C. SnapShot: directed differentiation of pluripotent stem cells. Cell 149, 1174–1174 e1171, doi:10.1016/j.cell.2012.05.015 (2012).

2. Clevers, H. Modeling Development and Disease with Organoids. Cell 165, 1586–1597, doi:10.1016/j.cell.2016.05.082 (2016).

3. Lancaster, M. A. & Knoblich, J. A. Organogenesis in a dish: modeling development and disease using organoid technologies. Science 345, 1247125, doi:10.1126/science.1247125 (2014).

4. Green, R. E. et al. A draft sequence of the Neandertal genome. Science 328, 710–722, doi:10.1126/science.1188021 (2010).

5. Prufer, K. et al. The complete genome sequence of a Neanderthal from the Altai Mountains. Nature 505, 43–49, doi:10.1038/nature12886 (2014).

6. Prufer, K. et al. A high-coverage Neandertal genome from Vindija Cave in Croatia. Science 358, 655–658, doi:10.1126/science.aao1887 (2017).

7. Vernot, B. & Akey, J. M. Resurrecting surviving Neandertal lineages from modern human genomes. Science 343, 1017–1021, doi:10.1126/science.1245938 (2014).

8. Simonti, C. N. et al. The phenotypic legacy of admixture between modern humans and Neandertals. Science 351, 737–741, doi:10.1126/science.aad2149 (2016).

9. Khrameeva, E. E. et al. Neanderthal ancestry drives evolution of lipid catabolism in contemporary Europeans. Nat Commun 5, 3584, doi:10.1038/ncomms4584 (2014).

10. Sankararaman, S. et al. The genomic landscape of Neanderthal ancestry in present-day humans. Nature 507, 354–357, doi:10.1038/nature12961 (2014).

11. Quach, H. et al. Genetic Adaptation and Neandertal Admixture Shaped the Immune System of Human Populations. Cell 167, 643–656 e617, doi:10.1016/j.cell.2016.09.024 (2016).

12. Dannemann, M. & Kelso, J. The Contribution of Neanderthals to Phenotypic Variation in Modern Humans. American journal of human genetics 101, 578–589, doi:10.1016/j.ajhg.2017.09.010 (2017).

13. Sams, A. J. et al. Adaptively introgressed Neandertal haplotype at the OAS locus functionally impacts innate immune responses in humans. Genome biology 17, 246, doi:10.1186/s13059-016-1098-6 (2016).

14. Dannemann, M., Andres, A. M. & Kelso, J. Introgression of Neandertal – and Denisovan-like Haplotypes Contributes to Adaptive Variation in Human Tolllike Receptors. American journal of human genetics 98, 22–33, doi:10.1016/j.ajhg.2015.11.015 (2016).

15. Consortium, S. T. D. et al. Sequence variants in SLC16A11 are a common risk factor for type 2 diabetes in Mexico. Nature 506, 97–101, doi:10.1038/nature12828 (2014).

16. McCoy, R. C., Wakefield, J. & Akey, J. M. Impacts of Neanderthal-Introgressed Sequences on the Landscape of Human Gene Expression. Cell 168, 916–927 e912, doi:10.1016/j.cell.2017.01.038 (2017).

17. Dannemann, M., Prufer, K. & Kelso, J. Functional implications of Neandertal introgression in modern humans. Genome biology 18, 61, doi:10.1186/s13059-017-1181-7 (2017).

18. Kilpinen, H. et al. Common genetic variation drives molecular heterogeneity in human iPSCs. Nature 546, 370–375, doi:10.1038/nature22403 (2017).

19. Genomes Project, C. et al. A global reference for human genetic variation. Nature 526, 68–74, doi:10.1038/nature15393 (2015).

20. Meyer, M. et al. A high-coverage genome sequence from an archaic Denisovan individual. Science 338, 222–226, doi:10.1126/science.1224344 (2012).

21. Sankararaman, S., Mallick, S., Patterson, N. & Reich, D. The Combined Landscape of Denisovan and Neanderthal Ancestry in Present-Day Humans. Current biology: CB 26, 1241–1247, doi:10.1016/j.cub.2016.03.037 (2016).

22. Vernot, B. et al. Excavating Neandertal and Denisovan DNA from the genomes of Melanesian individuals. Science 352, 235–239, doi:10.1126/science.aad9416 (2016).

23. Qin, P. & Stoneking, M. Denisovan Ancestry in East Eurasian and Native American Populations. Molecular biology and evolution 32, 2665–2674, doi:10.1093/molbev/msv141 (2015).

24. Huerta-Sanchez, E. et al. Altitude adaptation in Tibetans caused by introgression of Denisovan-like DNA. Nature 512, 194–197, doi:10.1038/nature13408 (2014).

25. Kong, A. et al. Fine-scale recombination rate differences between sexes, populations and individuals. Nature 467, 1099–1103, doi:10.1038/nature09525 (2010).

26. Hinch, A. G. et al. The landscape of recombination in African Americans. Nature 476, 170–175, doi:10.1038/nature10336 (2011).

27. Zerbino, D. R., Wilder, S. P., Johnson, N., Juettemann, T. & Flicek, P. R. The ensembl regulatory build. Genome biology 16, 56, doi:10.1186/s13059-015-0621-5 (2015).

28. Consortium, G. T. et al. Genetic effects on gene expression across human tissues. Nature 550, 204–213, doi:10.1038/nature24277 (2017).

29. Danecek, P. et al. The variant call format and VCFtools. Bioinformatics 27, 2156–2158, doi:10.1093/bioinformatics/btr330 (2011).

